# Rapid multiplex small DNA sequencing on the MinION nanopore sequencing platform

**DOI:** 10.1101/257196

**Authors:** Shan Wei, Zachary R. Weiss, Zev Williams

## Abstract

Real-time sequencing of short DNA reads has a wide variety of clinical and research applications including screening for mutations, target sequences and aneuploidy. We recently demonstrated that MinION, a nanopore-based DNA sequencing device the size of a USB drive, could be used for short-read DNA sequencing. In this study, an ultra-rapid multiplex library preparation and sequencing method for the MinION is presented and applied to accurately test normal diploid and aneuploidy samples’ genomic DNA in under three hours, including library preparation and sequencing. This novel method shows great promise as a clinical diagnostic test for applications requiring rapid short-read DNA sequencing.

## Introduction

Rapid sequencing of short DNA reads may be useful for a wide range of clinical and-research applications including targeted mutation analysis, cancer-panel testing, and aneuploidy screening (Kukita *et al.* 2015; Zheng *et al.* 2015; Butler *et al.* 2016). However, the time and skill required for library preparation and sequencing using existing DNA sequencing methods limits their widespread clinical use. Nanopore sequencing technology is one of the fastest growing 3^rd^ generation next-generation sequencing (NGS) technologies (Haque *et al.* 2013; Wang *et al.* 2014; Jain *et al.* 2016). Different from the 2^nd^ generation NGS sequencing platforms, such as illumina MiSeq and Ion Proton, the 3^rd^ generation NGS platforms, including nanopore sequencing, sequenced nucleotides at single-molecule level (Quail *et al.* 2012; Jain *et al.* 2016). MinION was the first portable nanopore sequencing platform to be commercially released (Loman and Watson 2015). It detects the electric current of single-stranded DNA (ssDNA) as it passes through a small protein channel, called a nanopore, and converts the electric current data into the corresponding sequence (Loman and Watson 2015). As this method relies on the physicochemical properties of ssDNA rather than an enzymatic reaction, sequencing occurs at speeds that are faster than 2^nd^ generation NGS systems (15,000 nt/min vs. 1 nt/min using Ion Proton).

While MinION was originally developed for sequencing long strands of DNA (>8 kb and even >1,000 kb), it was recently demonstrated that changes in the chemistry and library preparation could allow the device to be used for sequencing of short DNA reads (∼500 nt) (Wei and Williams 2016). However, library preparation for a single sample took 4 hours and multiple technically complex steps to complete, sequencing took an additional 1-4 hours, and only a single sample could be sequenced at a time (Figure 1). These factors limited the clinical utility of the MinION.

In this study, we report a new method that simplifies and accelerates nanopore-based short-length DNA library preparation and sequencing, and apply this method to test a panel of normal and aneuploid genomic DNA samples. We were able to accurately screen for aneuploidy using pure genomic DNA in 5 multiplexed samples in under 3 hours, including the time required for library preparation and sequencing. Using this method, the MinION nanopore sequencer can be used for a wide range of applications requiring rapid short-read DNA sequencing.

**Figure 1.**
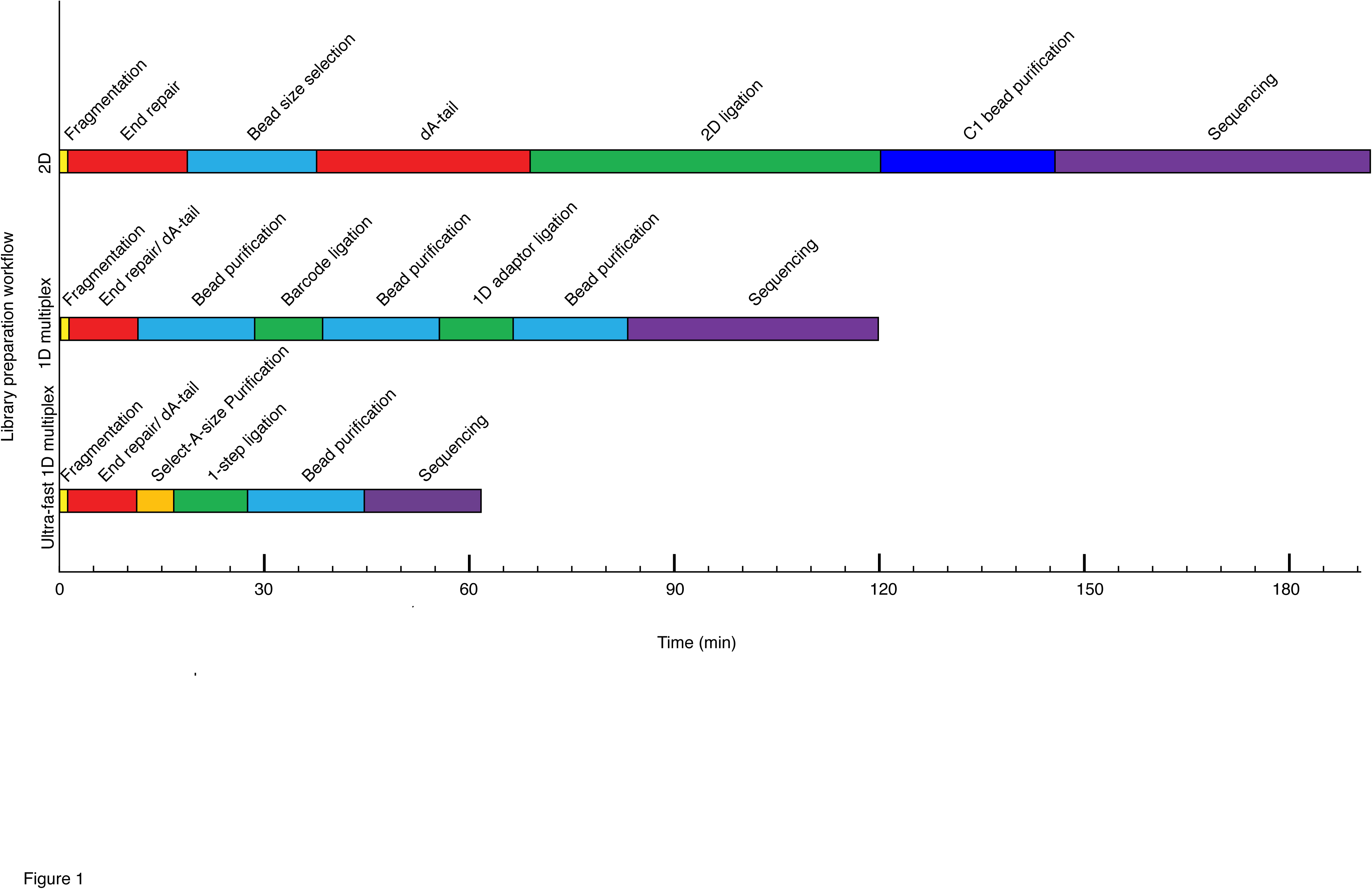
Comparison of MinION library preparation workflows. 2D library is a previously reported workflow (Wei and Williams 2016); 1D multiplex library is manufacturer’s workflow using a native barcoding kit and a 1D genomic sequencing kit on the current MinION platform; Rapid 1D multiplex library is a new rapid barcoding MinION library preparation workflow reported in this study developed to sequence short reads (<1000 bp) on the current platform. The length of each bar indicates the time needed. The steps are color-coded (Yellow: fragmentation; Red: end preparation including end-repair and dA-tail; blue: size selection and purification; dark blue: MyOne C1 bead purification; purple: sequencing)

## Materials and Methods

### DNA Samples

Normal male (NA12877), normal female (NA12878), Monosomy X (NG08006), trisomy 21 (NG05397) and trisomy 18, 15s+ (NG08016) genomic DNA (gDNA) from the Coriell Institute were used for development and testing of this protocol. The study was approved by the Institutional Review Board of Albert Einstein College of Medicine and the Institutional Review Board of Columbia University Medical Center and complied with Coriell Institute NIGMS Human Genetic Cell Repository and NIA Aging Cell Culture Repository policies.

### Development of rapid ligation conditions

To develop conditions for rapid ligation that could combine ligation of both the TA end and the 6-bp sticky-end, thus reducing the time needed for library preparation, a range of ligation enhancers were tested. The ligation efficiency of 6-bp sticky-end ligation and TA ligation was estimated using different adapters for each ligation. For the 6-bp sticky-end ligation, 2 pmol of a 58 bp asymmetric adapter with a 3’-ATTGCT overhang (MP1-6bp) and 2 pmol of a 34 bp adapter with a 3’-AGCAAT and 5’ blunt-end (ME-6bp) were used (Table S1) (Wei and Williams 2016). In addition to the 10 μL basic ligation substrates (2 μL of 1 μM MP1-6bp adapter, 2 μL of 1 μM ME-6bp adapter, 6 μL of 2 mM Tris-Cl, pH 8 (1/5 Buffer EB, Qiagen, Cat. 19086)), 50 or 10 μL Blunt/TA ligase master mix (NEB, Cat. M0367) and 0, 1.2, 1.8 or 2.4 μL enhancer mix (83.3 mM MgCl_2_, 16.7 mM ATP) were added to each ligation reaction and incubated at 25°C for 10 min (Figure 2A). Each ligation reaction was purified using a DNA Clean & Concentrator-5 column (Zymo, Cat. D4003 or D4013) following the manufacturer’s protocol. Each reaction mixture was mixed with DNA binding buffer at a 1:7 volume ratio (Zymo, Cat. D4003 or D4013); the ligation products were eluted in 20uL 2 mM Tris-Cl, pH 8, and products ≥ 30 bp were retrieved. The purified ligation products were then analyzed using 3% agarose gel electrophoresis and ImageJ (imagej.nih.gov/ij/) densitometry analysis with 2 technical replicates (Figure 2A).

**Figure 2.**
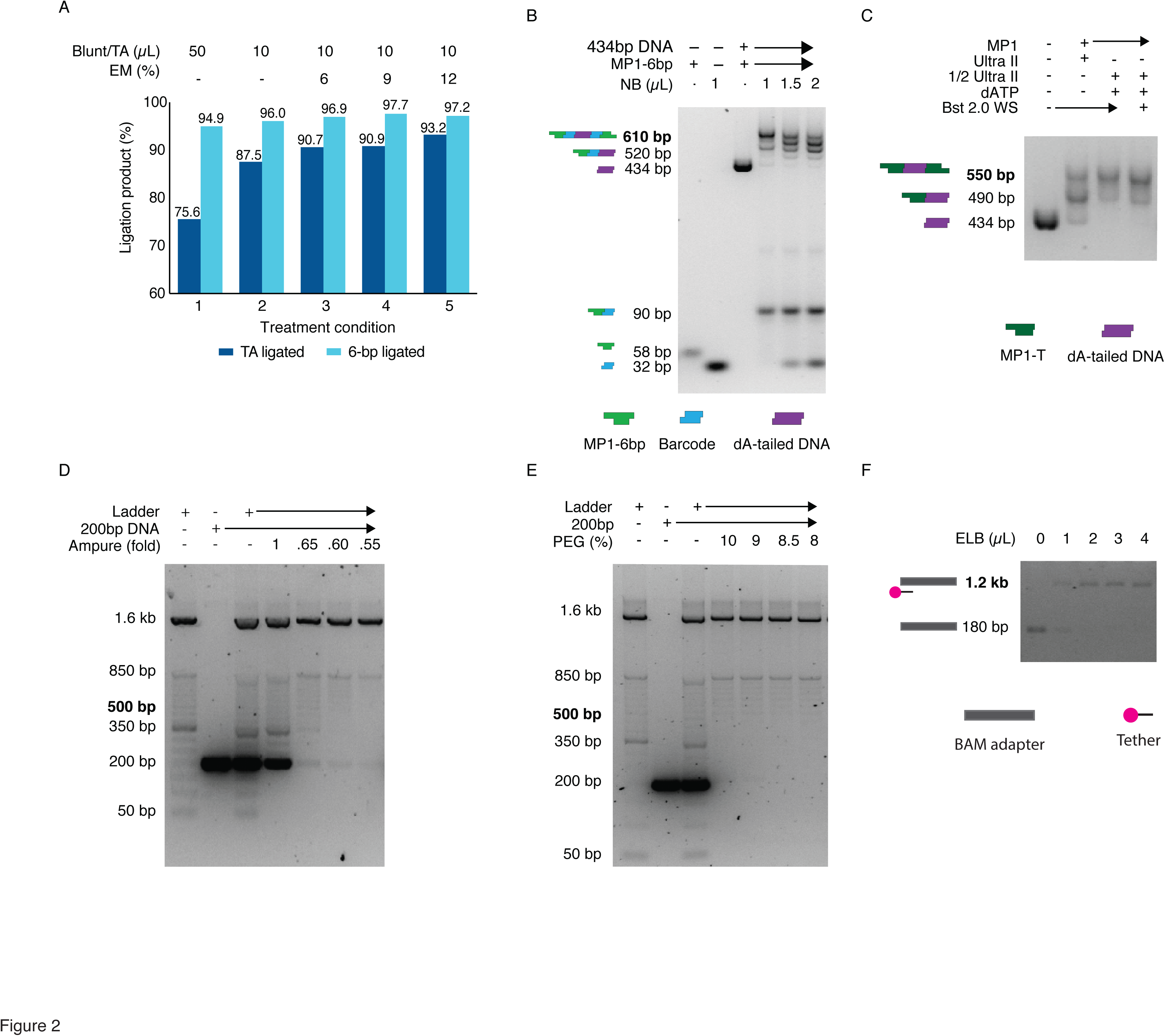
Optimization of MinION library preparation. A). Optimization of ligation condition for TA ligation and 6-bp sticky-end ligation. Condition 1. The manufacturer’s suggested condition; 2. the condition reported before (Wei and Williams 2016); 3-5. the conditions with addition of 6%, 9%, and 12% enhancer mix. Efficiencies of 6-bp ligation were estimated using a pair of adaptor MP1-6bp and ME-6bp carrying complementary 6-bp sticky ends. Efficiencies of TA ligation were estimated using a pair of adaptor MP1-T and ME-A carrying complementary 3’T and 3’A overhangs. B). Titration experiment of Native Barcode (NB) adapter. 6.5ng, 9.8ng, 13ng of NB adapters were added in to the 1-step ligation reaction which contains the same amount of dA-tailed DNA and MP1-6bp adapter. The expected final products with 2-end ligated to a barcode and MP1-6bp adapter were marked in bold. The products separated on gel were also illustrated in cartoons (MP1-6bp adapter: green; NB adapter: blue; dA-tailed DNA: purple). C). Optimization of end-repair/dA-tailling condition. Lane 1, the input 434bp control fragment; lane 2, manufacturer’s recommended protocol; lane 3. the optimized condition; lane 4. The optimized condition with supplementation of Bst 2.0 WarmStart Polymerase. The expected products with 2-end ligated to an adapter were marked in bold and the products separated on gel were also illustrated in cartoons (434bp dA-tailed DNA: purple; MP1-T adapter: dark green). D). Optimization of AMPure XP bead purification by changing the volume of bead. 100 ng 50bp ladder and 2 pmole 204bp control fragment were used as input, and subjected to 1-fold, 0.65-fold, 0.6-fold, 0.55-fold AMPure XP bead purification. The expected products are bands >500 bp, and it’s marked in bold E). Optimization of AMPure XP bead purification by adjusting the concentration of PEG in wash buffer. 100 ng 50bp ladder and 2 pmole 204bp control fragment were used as input, and subjected to 0.62-fold AMPure XP bead purification using wash buffer containing 10%, 9%, 8.5% and 8% PEG. The expected products are bands >500 bp, and it’s marked in bold F). Optimization of tethering condition. Lane 1-5: 1μL BAM adapter with 0-4μL ELB buffer after 3min incubation at 37°C. The expected tethered BAM adapter was marked in bold. The products separated on gel were illustrated in cartoons (BAM adapter: gray; tether: pink-black).

The efficiency of the TA ligation was estimated by the same method as 6-bp sticky-end ligation but using a different pair of adapters that had 3’ T/A overhangs (MP1-T and ME-A) (Table S1). 4uL 200 nM MP1, 4 μL 200 nM ME-A adapters, and 2 μL 2 mM Tris-Cl, pH 8 were subjected to each ligation condition as described above (Figure 2A).

Although the 12 native barcoding (NB) adapters in a 1D native barcoding kit (Oxford Nanopore, EXP-NBD103) are designed to be supplied at 670 nM (communications with the manufacturer’s technical support), the actual concentration of each NB adapters in a kit can vary from 2-13 ng/μL. Thus, the DNA content of each NB adapter was quantified using the Qubit dsDNA HS assay (Invitrogen, Cat. Q32851). To determine the right amount of NB adapter needed in a 1-step ligation reaction, a titration ligation experiment was performed. 6.5 ng, 9.8 ng, or 13 ng NB adapters were added to a 21.2 μL ligation reaction mixture containing 0.2 pmole dA-tailed 434bp, 1.6 pmole MP1-6bp, 10 μL Blunt/TA ligase master mix, and 1.2 μL enhancer mix (83.3 mM MgCl2, 16.7 mM ATP) in 2 mM Tris-HCl. Each ligation reaction mixture was incubated at 25 °C for 10 min, and purified using a DNA Clean & Concentrator-5 column as described above. The purified ligation products were analyzed using 3% agarose gel electrophoresis and ImageJ densitometry analysis (imagej.nih.gov/ij/) (Figure 2B).

### Development of rapid end-repairing/dA tailing condition

A short DNA control fragment was used to assess the efficiency of different conditions for end-repairing/dA-tailing. This fragment was generated by PCR using M13 forward and reverse primers to amplify a 434 bp fragment from a *pCR-Blunt* vector (Invitrogen, Cat. K2700-20) using Q5^®^ High-Fidelity DNA Polymerase (NEB, Cat. M0491S) (Table S1) as reported previously (Williams and Wei, 2016).

The resulting fragment was mixed with the NEBNext^®^ Ultra^™^ II End Repair/dA-Tailing Module (NEB, Cat. E7546L) in a clean 8-well 0.2 mL PCR strip and incubated for 5min at 20°C for end-repair reaction, and then 5 min at 65°C for dA-tailing reaction. 3 conditions were tested to optimize end-repair/dA-tailing: 1) ∼0.5 pmol DNA (∼162.5 ng) in 60uL total reaction mixture; 2) ∼0.5 pmol DNA in 30 μL total reaction mixture with addition of 0.9 μL 100 mM dATP before the 65°C incubation; 3) ∼0.5 pmol DNA control fragment in 30 μL reaction mixture with addition of 0.9 μL 100 mM dATP and 1μL Bst 2.0 WarmStart DNA polymerase (NEB, Cat. M0538S) before the 65°C incubation. Each reaction mixture was then purified using a DNA Clean & Concentrator-5 column (Zymo, Cat. D4003 or D4013) following the manufacturer’s protocol as described above.

The efficiency of end-repair/dA-tailing of each of the 3 conditions was assessed using a 57-bp asymmetric adapter with a 3’T overhang (MP1-T) (Table S1). The adapter was diluted to 0.4 μM in MinION adapter buffer (50 mM NaCl, 10 mM Tris-HCl, pH 7.5), and ligated to the dA-tailed control fragment in a 10:1 ratio as previously reported (Wei and Williams 2016). 21.2 μL ligation reaction included 4 μL 50 nM dA-tailed DNA, 5 μL 0.4 μM adapter, 1 μL nuclease-free water (Ambion, Cat. AM9937), 10 μL Blunt/TA ligase master mix (NEB, Cat. M0367S), and 1.2 μL enhancer mix (83.3 mM MgCl2, 16.7 mM ATP). The ligation reaction was incubated at 25°C for 10 min, purified using a DNA Clean & Concentrator-5 column as above, and analyzed with 3% agarose gel electrophoresis and ImageJ densitometry analysis (imagej.nih.gov/ij/) (Figure 2C).

### Development of library purification conditions

10% PEG 8000 (PEG) wash buffer (10% PEG, 750 mM NaCl, 50 mM Tris-HCl, pH=8) was prepared following the protocol in the Oxford Nanopore MAP003 kit. 9%, 8.5%, and 8% PEG buffers were also prepared to determine which concentration would most efficiently remove excessive adapters through AMPure XP bead purification (Agencourt, Cat. A63881). PEG wash buffers were stored at 4°C. 2 pmole (∼260 ng) of a 204 bp DNA fragment (Table S1) and 100 ng of a 50 bp ladder (Invitrogen, Cat. 10416014) were mixed in Buffer EB (Qiagen, Cat. 19086) to make a 60 μL reaction mixture.

To determine the optimal concentration of AMPure XP beads for the 1-step ligation reaction purification, beads were added to the reaction mixture at 1:1, 0.65:1, 0.60:1, or 0.55:1 bead: sample ratios in each well of a 0.2 mL PCR strip (USA Scientific, Cat. 1402-4700) and purified according to the manufacturer’s protocol on a 96-well magnet plate (ALPAQUA, Cat. A001322). However, in the wash steps, the 10% PEG wash buffer described above was substituted for the 70% ethanol used in the manufacturer’s protocol. After two washes, the beads were resuspended in 20 μL Buffer EB (Qiagen, Cat. 19086) and incubated at 37°C for 5 min. The resuspended beads were allowed to pellet on the magnet plate, and the eluate was carefully transferred to a 1.5 mL low retention tube (USA Scientific, Cat. 1415-2600 or Phenix, Cat. MH-815S). The AMPure XP bead purified products were analyzed with 2% agarose gel electrophoresis (Figure 2D).

To determine the optimal PEG concentration in PEG wash buffer to remove extra adapters, AMPure XP bead purification was performed using a 0.62:1 beads: sample ratio and washed in each PEG wash buffer described above (Figure 2E).

To test the efficiency of adapter removal for multiple reactions, 8 reaction mixtures were purified by 0.62X AMPure XP bead purification and washed in 8% PEG buffer as described above. The efficiency was not altered (data not shown).

### Development of library tethering conditions

At the end of library preparation, each 1D barcoding sequencing adapter (BAM) (Oxford nanopore, EXP-NBD103) must be annealed to tethering oligonucleotides (tethers), which carry a hydrophobic group on their 5’end, included in the elution buffer (ELB) (Oxford nanopore, SQK-LSK108). This process is called tethering in the manufacturer’s protocol. Tethering in ELB buffer can assist the barcode sequencing adapters (BAM) to reach the nanopores faster. Motor proteins are pre-attached to BAM adapters. When a BAM adapter reaches a nanopore, the motor protein can unzip dsDNA into ssDNA and drive the resulting DNA strand through the nanopore at a fixed speed. The ideal tethering condition was determined by mixing the BAM adapter with ELB buffer in 1:1, 1:2, 1:3, and 1:4 ratios and incubated at 37°C for 3min. The BAM adapter was completely tethered at a 1:2 BAM: ELB ratio (Figure 2F). Tethering was also tested at 25°C and on ice at a 1:2 ratio for 10min but these conditions were less efficient than tethering at 37°C for 3min (data not shown).

### Rapid multiplex MinION Sequencing library preparation

Each optimized step was combined to form our rapid multiplex MinION sequencing library preparation. ∼500 ng gDNA was fragmented with a Covaris microTUBE using the manufacturer’s default 500-bp setting. ∼0.5 pmole of fragmented gDNA (165 ng ∼500-bp DNA) was subjected to a 30 μL end-repair/dA-tailing reaction using the NEBNext^®^ Ultra^™^ II End Repair/dA-Tailing Module (NEB, Cat. E7546L) as detailed above with addition of 3 mM dATP after 5min incubation at 20°C, followed by 5 min incubation at 65 °C. A Select-a-Size DNA Clean & Concentrator (Zymo, Cat. D4080) was used to purify each end-repair/dA-tail reaction following the manufacturer’s protocol. 156 μL size selection mix (500 μL Select-a-Size DNA binding buffer + 5 μL 100% ethanol, can be scaled up for more reactions) was added to each end-repair/dA-tailing reaction tube and gently mixed by pipetting. The size-selected DNA was then eluted in 12 μL 2 mM Tris-Cl, pH 8.

A 1D native barcoding kit (Oxford Nanopore, EXP-NBD103) and a 1D Ligation Sequencing Kit (Oxford Nanopore, SQK-LSK108) were used for library preparation. As the native barcoding (NB) adapters in each kit (Oxford Nanopore, EXP-NBD103) can vary in concentration, the DNA content was quantified using the Qubit dsDNA HS assay (Invitrogen, Cat. Q32851). 6-10 ng NB adapter were used for each adapter ligation reaction.

In each 1-step ligation reaction, 15 μL ligation components (65 ng size selected dA-tailed DNA, 6-10 ng NB adapter, 8 μL BAM adapter, topped up to 15 μL using 2 mM Tris-HCl), 15 μL Blunt/TA ligase master mix, and 3.6 μL enhancer mix were mixed thoroughly. The mixture was then incubated at 25°C for 10 min. Two or more ligation reactions can be performed for 1 sample using 1 barcode to increase the final library concentration. In run 1, 1 NB adapter was used to barcode 1 sample and 4 ligation reactions were prepared for this sample; In run 2 and run 3, 5 NB adapters were used to barcode 5 sample and 2 ligation reactions were used for each sample (Table 1). 26.4 μL Buffer EB and 21.6 μL AMPure XP beads were added to each ligation reaction to perform bead purification on a magnet plate (equivalent to ∼0.62 fold beads: sample ratio when counting the PEG in Blunt/TA ligase master mix). 8% PEG wash buffer was used as described above. The beads from all the ligation reactions were pooled in a clean 1.5 mL low retention tube after the first wash, pelleted, and washed again on a magnet stand (Agencourt, Cat. A29182). The resulting library was eluted in 16-20 μL ELB buffer by incubation at 37°C for 3-5 min. The DNA concentration was quantified with Qubit dsDNA HS assay using 1μL library. The final concentration of the library was ∼15-20 ng/μL. In manufacturer’s protocol, for DNA shorter then 3kb, it’s suggested to use 0.2 pmoles gDNA for library preparation. The concentration would be ∼ 6ng/μL for a library for 500bp fragments. It’s not sufficient to generate sufficient yields. It’s suggested to use a pre-sequencing library with ∼ 20ng/μL or higher to produce comparable results.

### MinION sequencing

12 μL of the library was loaded into a MinION MIN106 flow cell for sequencing following manufacturer’s protocol. MINKnow v1.3.30 software and run protocol NC_48Hr_sequencing_Run_FLO-MIN106_SQK-LSK108_plus_Basecaller were used to control and monitor the sequencing in real time. To maximize the data output for rapid MinION sequencing, the data acquisition was restarted after a 1 h run, and stopped when sufficient data was generated. A 30 min sequencing run generated enough reads for 1 sample; a 1-3 h sequencing run generated enough reads for 5 barcoded samples. The average number of real-time pores used for strand sequencing was 114-145 as monitored on the MINKnow software and the total number of pores generated sequences was >500 (data not shown).

### Data analysis

The MinKNOW run protocol NC_48Hr_sequencing_Run_FLO-MIN106_SQK-LSK108_plus_Basecaller generated sequencing results in FAST5 format. Sequences that passed the protocol’s quality filter were converted from FAST5 format to FASTA format using Poretools version 0.51 (Loman and Quinlan 2014; Wei and Williams 2016). The FASTA files were processed with Cutadapt version 1.14 to demultiplex the data using parameter (-O 20–e 0.20–m 50) (Martin 2011). At least a 20 bp match to a barcode with a maximum error rate of 0.20 was required to be considered a match to the barcode. Sequences ≥ 50 bp after demultiplexing were kept for downstream analysis. Sequences were aligned to human reference genome GRCh37 using Pblat (BLAT in parallel setting) (icebert.github.io/pblat/) using the parameters stated in Supplementary Table 2 (Table S2) (Kent 2002; Wei and Williams 2016). The performance of Pblat was evaluated using 20,000 sequences sampled from run 2 (Table S2). A new alignment software, minimap2, was also evaluated (Table S2) (Li 2017). This software is designed for various platforms including nanopore sequencing. It requires less computational resources and less run time at the cost of 5-6% less reads that can be used for downstream analysis. In this study, we used the alignment results from Pblat software to achieve the maximum number of reads for downstream data analysis.

The first 9,000 uniquely assigned (UA) from each sample were used for aneuploidy detection analysis. Given 9,000 UA reads, > 41 UA reads were assigned to chrY in a normal male sample. Data analysis and statistical analysis were performed in R version 3.4.0 (Team 2012). Under Poisson distribution, when 41 UA reads were assigned to one chromosome (λ = 41), the type I error for false positive detection of a 50% increase (e.g., a full trisomy) p(x > 1.5 λ) = 0.0008; the type II error for false negatively detecting a 50% changes p_β_(x’ < 1.5 λ) = 0.0021 and the type II error for false negatively detecting a 30% changes p_β_(x’ < 1.3 λ) = 0.04 as estimated by ppois function in R. UA reads aligned to each chromosome were summarized and analyzed using the modified Z-score method to identify normal diploid and aneuploid chromosomes as reported previously (Wei and Williams 2016). A known normal male sample was used as a reference. The standard deviation of the relative chromosomal copy number of normal autosomes included in this study, sd_normal = 0.0897 (n=219), was used to determine aneuploid chromosomes using the modified Z-score method as reported before (Table S3) (Wei and Williams 2016).

### Results

In this study we developed a rapid and practical protocol for preparing and sequencing a 1D multiplex genomic DNA short-read sequencing library using a nanopore sequencer. This was achieved by systematically and quantitatively evaluating and optimizing each module in the library preparation workflow. This provided robust reaction conditions in each module, and reduced required library preparation time from 2-4 h to ∼45 min and sequencing time from 1-4 h to < 30 min when compared to the original protocol (Figure 1) (Wei and Williams 2016).

A 2D library generates sequences from both template and complementary strand of each DNA fragment; a 1D library generated sequence from the template strand (Jain *et al.* 2016). On previous MinION MAP006 platform, a 2D library was prepared to increase the sequencing accuracy (Figure 1). However, on current MinION MIN106 platform, the sequencing quality was improved due to improvements on nanopore protein and basecalling algorithm (Jain *et al.* 2015; Carter and Hussain 2017), a 1D library can be prepared to generate sequences of comparable sequence quality as a 2D library on old platform (Figure 1).

A rapid 1D multiplex library includes 5 steps (Figure 1). 1) DNA fragmentation: DNA is sheared to ∼500bp by ultrasonication. 2). End preparation: sheared DNA fragments are repaired to blunt-ends with 5’ phosphorylation modification, and then 3’dA-tails are added. 3). Size selection to remove reads < 400bp: Zymo Select-a-Size columns can perform size selection during column purification to remove reads <400 bp. 4) 1-step ligation for barcodes and sequencing adapters: a native barcode (NB) adapter carries a 3’T overhang on one end that can be ligated to a 3’dA-tailed DNA fragment; it carries a 3’ 6bp overhang on the other end that can be ligated to the 1D barcode sequencing adapter (BAM). The TA ligation between DNA fragments and NB adapters and the 6-bp sticky-end ligation between NB and BAM adapters are carried out in one optimized ligation reaction at the same time. 5) Purification: the ligation reaction needs to be purified to retrieve fragments that are attached to NB and BAM adapters and eliminate unnecessary materials such as enzyme, extra adapters, etc. There is a pre-attached motor protein on BAM adapters which will assemble on a nanopore to unzip dsDNA into ssDNA strands and drive it for nanopore sequencing (Jain *et al.* 2016). The motor protein is sensitive to protein denaturants, freeze-and-thaw cycles and heat. Hence a wash buffer containing PEG was used instead of 70% ethanol in washing steps during bead purification. When the library is eluted off the beads, there is a hidden tethering step to anneal tethering oligonucleotides in elution buffer (ELB) to BAM adapters on fragments. The tethering oligonucleotides can help the fragment to reach a nanopore faster. This is done simultaneously at the end of the library purification step and no extra action needs to be taken. The rapid 1D barcoding library preparation method we present is a simplified and optimized method comparing to the current manufacturer’s 1D native barcoding library preparation method.

### Development of rapid ligation conditions

Purifications are the most time-consuming steps in a library preparation workflow. It can add 15-30 min to each step during library preparation (Figure 1). To bypass several ligation/purification steps in manufacturer’s 1D native barcoding library preparation workflow (Figure 1), we sought to develop a 1-step ligation condition that could allow simultaneous ligation for TA and 6-bp sticky ends, thereby enabling dA-tailed fragments to be ligated to a native barcoding (NB) adapter and then to a barcoding sequencing adapter (BAM) in one step. The NB adapter is designed asymmetrically with a 3’ 6-bp on one end and a 3’ T overhang on the other end. To quantitatively evaluate the ligation efficiency of 6bp sticky end ligation, a 62-bp asymmetric adapter (MP1-6bp) (Table S1) with a 6bp overhang and a 34 bp adapter (ME-6bp) (Table S1) were mixed with the complementary 6-bp overhang in a 1:1 ratio and subjected to a range of ligation conditions (Figure 2A). The portions of ligated and unligated MP1-6bp adapters were analyzed by 3% agarose gel electrophoresis and measured by densitometric analysis. To evaluate the corresponding ligation efficiency of TA ligation, a 57-bp asymmetric adapter (MP1-T) (Table S1) was mixed with a 3’T overhang and a 20 bp adapter (ME-A) (Table S1) was mixed with the complementary 3’A overhang in a 1:1 ratio and both mixtures were subjected to the ligation conditions described above (Figure 2A). Addition of enhancer mix increased the concentration of MgCl_2_ and ATP in a ligation system in a 5:1 ratio. Compared to the manufacturer’s protocol, using a 100 μL ligation mixture, 20 μL ligation conditions were more efficient on TA ligation, and robust on 6-bp sticky end ligation; unligated MP1-T adapter was reduced from 24.4% to 12.5%, and the unligated MP1-6bp adapter percentage from 5.1% to 4% (Figure 2A, Condition 2 vs. 1). In a 20 μL TA ligation reaction, addition of 6% to 12% volume of enhancer mix further reduced the percentage of unligated MP1 from 12.5% to 6.8-9.3%. Addition of 6% to 12% volume (1.2 to 2.4 μL) of enhancer mix further reduced the percentage of unligated MP1-6bp from 4% to ≤ 3% (Figure 2A). This method provides an efficient one-step ligation condition to simultaneously attach NB adapters to DNA fragments and BAM adapters to NB adapters without denaturing the motor protein of the BAM adapter.

During development, we noticed inconsistency in the concentration of NB adapters provided in the same kit. A titration experiment of NB adapter with a fixed amount of MP1-6bp adapter and dA-tailed DNA fragments was performed to determine the right range of NB adapter to be used in the 1-step ligation (Figure 2B). Adding 6.5 ng - 9.8 ng NB adapter resulted in most DNA fragments having NB adapters ligated on both sides, and most NB adapters were ligated. Only 1 BAM adapter is needed to sequence one DNA fragment in a 1D nanopore sequencing library. With addition of 13 ng NB adapters, NB adapters exceeded the amount of MP1-6bp adapter which resulted in more DNA fragments that ligated to NB adapters only but not MP1-6bp. In actual library preparation, ∼10ng NB for each ligation experiment was used as guided by this experiment.

### Development of rapid end-repairing/dA tailing condition

The NEBNext^®^ Ultra II End Repair/dA-Tailing Module (NEB, Cat. E7546L) combines end repair and dA-tailing in one reaction. We evaluated the efficiency of this product by mixing dA-tailed DNA samples with 10-fold diluted MP1 adapter (Table S1), and subjecting them to the optimized ligation method determined above. Using the same ligation conditions, TA ligation efficiency was affected by the proportion of fragments that had successfully been end-repaired and dA-tailed. The manufacturer’s Ultra II End-RepairR/dA-Tailing Module protocol did not provide comparable dA-tailing efficiency to that reported in our previous study (Figure 2C, lane 1) (Wei and Williams 2016), The majority of control fragments were 1-end ligated. However, reducing the volume of each preparation of the Ultra II End-Repair/dA-tailing module from 60 μL to 30 μL and supplying 0.9 μL 100 mM dATP after the 20°C incubation significantly improved the reaction efficiency from ∼30% two end-ligated to ∼80% (Figure 2C). During the development phase, we noticed batch effects on the Ultra II End-repair/dA-tailing module. Addition of 0.9 μL 100 mM dATP before the second incubation did not affect the ligation efficiency if the module had already reached its maximum ∼80% efficiency; it did, however, increase the efficiency to ∼80% if the module had only reached ∼50% efficiency (data not shown). Addition of more Bst 2.0 WarmStart DNA Polymerase (NEB, Cat. M0538) did not improve the dA-tailing efficiency (Figure 2C). The end-repairing/dA tailing could be performed in 12min, providing ∼80% 2-end ligation products, which is faster and more efficient than our previously reported system, which required > 1h for the reaction and purification, and provided ∼63% two-end ligated products (Wei and Williams 2016).

### Development of library purification conditions

There were two purification steps in the standard library preparation workflow. The first purification was performed after the end-repair/dA-tailing reaction. A Select-a-Size DNA Clean & Concentrator was used to perform a quick column-based reaction purification and size selection for short fragments < 400 bp. Addition of 5 μL 100% ethanol to 500 A Select-a-Size DNA binding buffer discards fragments < 400 bp while retaining fragments ≥ 400 bp efficiently (data not shown). The size selection and purification can be performed in ∼7 min.

The second purification was performed after the ligation reaction. This purification step is gentle to protein and efficient in removing excess adapters. AMPure XP beads with PEG-based wash buffer were used to protect the motor protein attached to the BAM sequencing adapter. We first used 0.62-fold equivalent AMPure XP beads to achieve size selection for fragments < 400 bp after the purification, but a noticeable amount of ∼200-bp adapter (BAM+NB) was retained after the bead purification. 1.2 to 2 pmol NB and BAM adapters were ligated to 0.2 pmol DNA fragments in a ligation reaction. To determine the optimal AMPure XP bead purification condition the purification efficiency was evaluated by mixing 100 ng 50 bp ladder and 2 pmol 204bp DNA fragment, and subjected to different purification conditions (Figure 2D, 2E). 1-fold AMPure XP bead did not eliminate most of the 200-bp DNA fragments. ≤ 0.65-fold AMPure XP bead reduced the 204-bp DNA fragment to <5% of the original input, but it was still visible when analyzed (Figure 2D). Reducing the AMPure XP volume to as low as 0.55-fold did not significantly reduce the 204-bp DNA fragment retention when compared to 0.65 and 0.60-fold AMPure XP bead. (Figure 2D). The wash step of the purification protocol was then optimized. The PEG-based wash in the MAP003 kit (Oxford nanopore, MAP003) contains 10% PEG, equivalent to the PEG concentration in a ∼0.8-fold AMPure XP bead purification reaction. The size selection effect in reducing the PEG concentration in PEG-based wash buffer from 10% to 9%, 8.5%, and 8% coupled with 0.65-fold AMPure XP bead purification was tested (Figure 2E). All conditions retained most of the fragments ≥ 450 bp. The wash buffers with 10%, 9% and 8.5% PEG retained visible 200 bp fragments, but no 200 bp fragments were visibly retained using 8% PEG wash buffer, the optimal condition.

### Development of library tethering conditions

To concentrate the oligonucleotide to be sequenced at the bottom of the sequencing flow cell where the nanopores are located, the BAM adapter needs to be tethered to an oligonucleotide in the ELB buffer that attached to a hydrophobic group. Using Dynabeads^™^ MyOne^™^ Streptavidin C1 bead-based 2D library purification (Invitrogen, Cat. 65001), the library was eluted from the beads in ELB buffer at 37 °C for 10 min (Figure 1) (Wei and Williams 2016). Using manufacture’s 1D library preparation protocol, the library was eluted from the beads in ELB buffer at RT °C for 10 min (Figure 1). This does not only elute the library from the beads, but also anneal the adapter to the tethering oligonucleotide. The tethering efficiency was tested by mixing 1 μL BAM adapter with 0-4 μL ELB buffer incubated at 37°C for 3 min (Figure 2E); it’s also tested by mixing 1 μL BAM adapter with 2 μL ELB buffer with incubation at RT for 10 min, and at 37 °C for 2 min, 5 min and 10 min (data not shown). 3-5 min incubation at 37 °C of 1 μL BAM and ≥ 2 μL ELB buffer was determined to be the optimal tethering condition.

### MinION sequencing and data analysis

Each of these optimized conditions combined to form a robust rapid multiplexed MinION Sequencing library preparation workflow in ∼45 min (Figure 1). The current MinION MIN106 platform provides higher sequencing quality for 1D reads than previous MinION platforms. Libraries were sequenced for 1-3 h and stopped when sufficient data was generated (Table 1). The read length of majority of reads felt between 500-1000bp, and the quality score (Q-score) of each base ranged from can range from 4 to 16 (Supple. Figure 1). The mean Q-score is ∼9-10 (Supple. Figure 1). The platform generated ∼70K raw reads per hour. 55-80% reads could be assigned to a unique barcode, and 92-95% reads with a barcode could be aligned to a unique genomic location (Figure 3A, Table 1). For a single sample, 12 min of sequencing generated 9K UA reads, 27 min generated 15K UA. This compares favorably to our prior method which required 57 min for the same number of reads (Table 1, Figure 3A) (Wei and Williams 2016). For barcoded samples, 1-3 h was sufficient to generate data for aneuploidy detection for up to 5 samples (Table 1). Three batches of samples were tested in this study, and aneuploidy was determined using the adjusted Z-score method reported in our previous study (Table 1, Table S3, Fig 3B-E) (Wei and Williams 2016). The normal male and female, trisomy and monosomy cases were detected concordantly with their karyotyping (Table 1).

This method does have limitations. During development, we evaluated the time needed using pure genomic DNA. In actual lab or clinical applications, time used to obtain sufficient DNA needs to be added. It takes ∼1h to extract gDNA from body fluids and cell lines, and 2-3h to extract gDNA from tissues, and longer for difficult samples such as fixed tissue and slides. Time and computational resources are also needed for data analysis. In the current pipeline, ∼ 30min is needed to analyze each sample (Table S2). Analyses can be performed in parallel in 30min given sufficient computational resources. A new nanopore sequence aligner, minimap2, can be a potential substitute in to perform data analysis in 10min with less computational resources at the cost of 5-6% loss in UA reads for downstream analysis (Table S2).

In this study, the enlargement of satellite DNA on chromosome 15 could not be detected due to the detection limit of ultra-low-coverage-sequencing (ULCS)-based methods (Table 1). ULCS-based method used sequence reads that can be uniquely aligned to human reference genome for data analysis; sequences aligned to highly repetitive regions such as satellite DNA and other genomic repetitive elements were eliminated from the analysis. The method is currently used for aneuploidy detection at the whole chromosome scale. At the current sequencing depth of 9K reads, we estimate that copy number variations (CNVs) can only be reliably detected if they are > 30MBs in length; to detect CNVs of <10Mb, we estimate that a read depth of >30K reads would be necessary. Reducing number of samples or increasing sequencing time might be needed for detection of large CNV events, though this would need to be confirmed experimentally.

**Figure 3.**
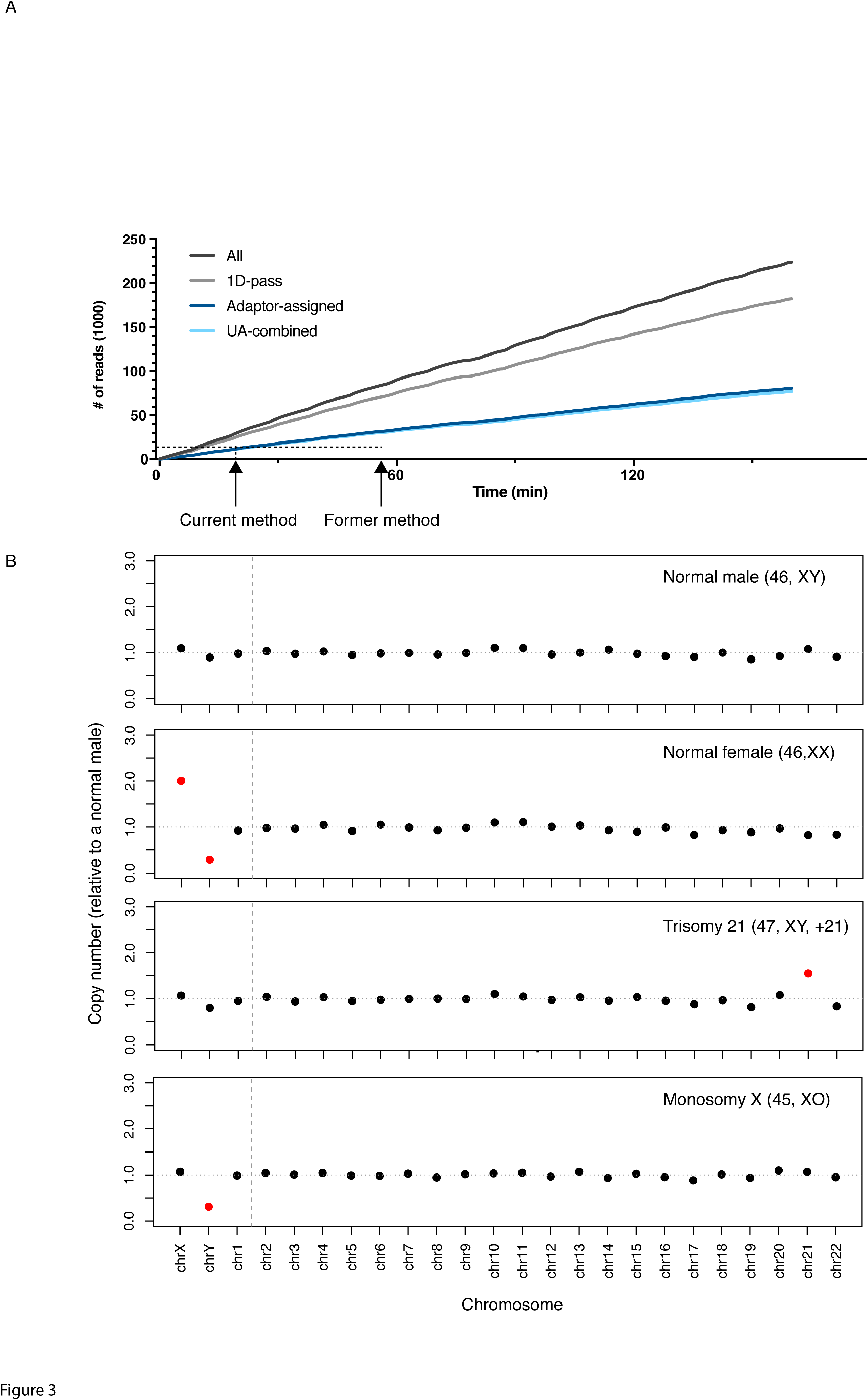
MinION Run performance and assay results. A). The run performance of the rapid 1D barcoding MinION library preparation method. B). Illustation of MinION assay results for a normal male, normal female, trisomy 21 and a monosomy X. Normal and aneuploidy on each chromosome was indicated by color. (Normal: black; Aneuploidy: red)

**Table 1.** MinION assay results

| Run | Barcode | Reads | Unique alignment | Time to reach 9K | MinION assay result | Sample karyotype |
| --- | --- | --- | --- | --- | --- | --- |
| 1 | B01 | 20,000* | 19637 | 0:12 | 46,XX | 46,XX |
| 2 | B02 | 21,583 | 20,171 | 0:54 | 45,XO | 45,XO |
| 2 | B03 | 18,320 | 17,143 | 1:05 | 47,XY,+21 | 47,XY,+21 |
| 2 | B04 | 18,501 | 17,647 | 1:03 | 46,XX | 46,XX |
| 2 | B05 | 18,998 | 17,975 | 1:00 | 46,XY | 46,XY |
| 2 | B08 | 20,406 | 19,097 | 0:56 | 47,XY,+18 | 47,XY,+18,15s+ |
| 3 | B01 | 19,184 | 18,347 | 1:09 | 46,XY | 46,XY |
| 3 | B02 | 9,272 | 8,818 | 2:42 | 46,XY | 46,XY |
| 3 | B03 | 12,033 | 11,535 | 2:01 | 46,XX | 46,XX |
| 3 | B05 | 18,626 | 17,762 | 1:15 | 47,XY,+21 | 47,XY,+21 |
| 3 | B06 | 21,793 | 20,823 | 1:02 | 45,XO | 45,XO |

\*40,045 reads with barcodes were generated in a 66min run. The first 20K reads with barcodes were subjected to downstream analysis

Last but not least, alignment-based copy number estimation as applied in this study has limitation in detecting some polyploid cases when the sex chromosomes are at the same ratio as a normal male or female (e.g., 69, XXX). A Karyotype or SNP-based approaches is more reliable in polyploidy detection at this point (Handyside 2011).

## Discussion

In this study, we developed a clinically viable protocol for rapid sequencing of short DNA fragments utilizing the MinION nanopore sequencer. As the nanopore sequencing platform was developed for ultra-long fragment sequencing, the manufacturer’s protocol is not ideal for preparing short-read sequencing libraries. Thus, we systematically optimized each module of the library preparation workflow, including the end repair/dA tailing reaction, column size selection, 1D barcoding ligation, and bead purification, to achieve a practical rapid 1D barcoded sequencing library preparation protocol. The adjustments in each step were critical to performing robust and efficient reactions and purifications. The resulting protocol is less dependent than previously described protocols on the version of the sequencing library kit used. It utilizes the NB and BAM adapters in the native barcoding kit and the ELB buffer in the 1D genomic sequencing kit that are readily available whenever a compatible version of flow cell is on the market.

We then successfully used this method for aneuploidy testing in genomic DNA samples. Using this method, up to five samples can be multiplexed to produce sufficient sequencing data for aneuploidy detection in 1-3h on a single flow cell, including library preparation time. It also takes additional 1-3h for gDNA extraction from human tissues or body fluids, and 30min for computation analysis in the development phase. Faster computation analysis can be achieved by allocating more computational resources and performing analysis on a SSD drive, or using newly developed software. Fetal aneuploidy testing is routinely performed as an essential component of prenatal testing (e.g. amniocentesis and chorionic villus sampling (CVS)), preimplantation genetic screening (PGS) of embryos after *in-vitro* fertilization (IVF), and evaluation of miscarriage tissue (Brezina *et al.* 2012; Wei and Williams 2016). A rapid diagnosis is clinically important as it enables timely management. Current standard methods to diagnose aneuploidy, such as karyotyping and microarray assays, take 7-21 days to complete (Reddy *et al.* 2012; Wapner *et al.* 2012; Dong *et al.* 2016). It also costs > $1,000 per assay. Ultra-low coverage sequencing (ULCS) for detection of aneuploidy is a new and powerful strategy for whole-genome aneuploidy detection with shorter turn-over time, but still requires 15-24 h to complete and requires technically advanced library preparation and costly sequencing platforms ($80,000-$128,000) that cannot be easily applied in a physician’s office or in low complexity settings (Chen *et al.* 2014; Dong *et al.* 2016; Wei and Williams 2016). Thus, the method described here is the fastest sequencing-based method for aneuploidy detection reported to date and testing can be performed for ≤ $150 per sample on an affordable sequencing platform ($1,000). It can be set up easily at the point-of-care or in low complexity settings.

## Conclusion

We reported a robust rapid multiplex short-read MinION sequencing library protocol for ultra-fast aneuploidy detection for single and multiple samples. It shows promise for translation to clinical applications with a low assay cost and the fastest turnover time to date.

## Acknowledgements

We sincerely thank Drs. Thomas Tuschl, Jan Vijg, Yousin Suh, Brynn Levy, Steven Josefowicz, Barry Coller and members of the Williams lab for their helpful input and advice with this project and manuscript. This research was supported by the National Institutes of Health Grant HD068546 and U19CA179564.

## Author contributions

Z.W. conceived of the project. S.W. performed the experiments and prepared figures. Z.W. and S.W developed the method and analyzed the data. Z. W., S.W., and Z.R.W. prepared the manuscript.

## Reference

Brezina, P. R., D. S. Brezina and W. G. Kearns, 2012 Preimplantation genetic testing. BMJ 345: e5908.

Butler, K. S., M. Y. L. Young, Z. Li, R. K. Elespuru and S. C. Wood, 2016 Performance characteristics of the AmpliSeq Cancer Hotspot panel v2 in combination with the Ion Torrent Next Generation Sequencing Personal Genome Machine. Regulatory Toxicology and Pharmacology 74:178–186.

Carter, J.-M., and S. Hussain, 2017 Robust long-read native DNA sequencing using the ONT CsgG Nanopore system. Wellcome open research 2: 23–23.

Chen, S., S. Li, W. Xie, X. Li, C. Zhang et al, 2014 Performance comparison between rapid sequencing platforms for ultra-low coverage sequencing strategy. PLoS One9:e92192.

Dong, Z., J. Zhang, P. Hu, H. Chen, J. Xu et al, 2016 Low-pass whole-genome sequencing in clinical cytogenetics: a validated approach. Genetics in Medicine 18: 940–948.

Handyside, A. H., 2011 PGD and aneuploidy screening for 24 chromosomes by genome-wide SNP analysis: seeing the wood and the trees. Reprod Biomed Online 23: 686–691.

Haque, F., J. Li, H.-C. Wu, X.-J. Liang and P. Guo, 2013 Solid-State and Biological Nanopore for Real-Time Sensing of Single Chemical and Sequencing of DNA. Nano today 8: 56–74.

Jain, M., I. T. Fiddes, K. H. Miga, H. E. Olsen, B. Paten et al, 2015 Improved data analysis for the MinION nanopore sequencer. Nat Methods 12: 351–356.

Jain, M., H. E. Olsen, B. Paten and M. Akeson, 2016 The Oxford Nanopore MinION: delivery of nanopore sequencing to the genomics community. Genome Biol 17:239.

Kent, W. J., 2002 BLAT–the BLAST-like alignment tool. Genome Res 12: 656–664.

Kukita, Y., R. Matoba, J. Uchida, T. Hamakawa, Y. Doki et al, 2015 High-fidelity target sequencing of individual molecules identified using barcode sequences: De novo detection and absolute quantitation of mutations in plasma cell-free DNA from cancer patients. DNA Research 22: 269–277.

Li, H., 2017 Minimap2: versatile pairwise alignment for nucleotide sequences.

Loman, N. J., and A. R. Quinlan, 2014 Poretools: a toolkit for analyzing nanopore sequence data. Bioinformatics 30: 3399–3401.

Loman, N. J., and M. Watson, 2015 Successful test launch for nanopore sequencing. Nature Methods 12: 303–304.

Martin, M., 2011 Cutadapt removes adapter sequences from high-throughput sequencing reads. EMBnetjournal 17:10–10.

Quail, M. A., M. Smith, P. Coupland, T. D. Otto, S. R. Harris et al, 2012 A tale of three next generation sequencing platforms: comparison of Ion Torrent, Pacific Biosciences and Illumina MiSeq sequencers. BMC genomics 13: 341–341.

Reddy, U. M., G. P. Page, G. R. Saade, R. M. Silver, V. R. Thorsten et al, 2012 Karyotype versus microarray testing for genetic abnormalities after stillbirth. N Engl J Med 367: 2185–2193.

Team, R. D. C., 2012 R: A language and environment for statistical computing. R Foundation for Statistical Computing, Vienna, Austria. ISBN 3-900051-07-0, URL www.R-project.org. R Foundation for Statistical Computing, Vienna, Austria.

Wang, Y., Q. Yang and Z. Wang, 2014 The evolution of nanopore sequencing. Front Genet 5: 449.

Wapner, R. J., C. L. Martin, B. Levy, B. C. Ballif, C. M. Eng et al, 2012 Chromosomal microarray versus karyotyping for prenatal diagnosis. N Engl J Med 367: 2175–2184.

Wei, S., and Z. Williams, 2016 Rapid Short-Read Sequencing and Aneuploidy Detection Using MinION Nanopore Technology. Genetics 202: 37–44.

Zheng, H., H. Jin, L. Liu, J. Liu and W.-H. Wang 2015 Application of next-generation sequencing for 24-chromosome aneuploidy screening of human preimplantation embryos. Molecular cytogenetics 8: 38–38.

